# Estimating cortical thickness trajectories in children across different scanners using transfer learning from normative models

**DOI:** 10.1101/2023.03.02.530742

**Authors:** C. Gaiser, P. Berthet, S.M. Kia, M. A. Frens, C. F. Beckmann, R. L. Muetzel, A. Marquand

## Abstract

This work illustrates the use of normative models in a longitudinal neuroimaging study of children aged 6-17 years and demonstrates how such models can be used to make meaningful comparisons in longitudinal studies, even when individuals are scanned with different scanners across successive study waves. More specifically, we first estimated a large-scale reference normative model using hierarchical Bayesian regression from N=40,435 individuals across the lifespan and from dozens of sites. We then transfer these models to a longitudinal developmental cohort (N=5,985) with three measurement waves acquired on two different scanners that were unseen during estimation of the reference models. We show that the use of normative models provides individual deviation scores that are independent of scanner effects and efficiently accommodate inter-site variations. Moreover, we provide empirical evidence to guide the optimization of sample size for the transfer of prior knowledge about the distribution of regional cortical thicknesses. We show that a transfer set containing as few as 25 samples per site can lead to good performance metrics on the test set. Finally, we demonstrate the clinical utility of this approach by showing that deviation scores obtained from the transferred normative models are able to detect and chart morphological heterogeneity in individuals born pre-term.

## 1. Introduction

Identifying structural or functional biomarkers of psychiatric and neurological illnesses across the lifespan has received increasing attention in recent years. Many of these disorders present symptoms that begin during childhood and adolescence (Bayer et al., 2021; Rogers & de Brito, 2016; Solmi et al., 2022; Whittle et al., 2020). There is however large inter-individual heterogeneity in symptoms and underlying biology (DeLisi, 2008; Fuhrmann et al., 2022; Mills et al., 2021; Tamnes et al., 2017), making it challenging to pinpoint the precise underlying neurobiological substrates. Longitudinal datasets provide particularly valuable insights on the temporal evolution of brain development and offer considerable potential to understand the emergence of psychopathology and to parse this heterogeneity across individuals.

To detect and understand this heterogeneity and atypicality, there is a need to better characterize typical neurodevelopment (Insel, 2014; Volpe, 2009). In recent years, the availability of large datasets has greatly assisted efforts to understand inter-individual variability in brain development (Bethlehem et al., 2022; Rutherford, Fraza, et al., 2022). For example, large scale studies using cortical volume, cortical thickness (CT) and surface area have identified a general decrease in these metrics with age, after adolescence (Bethlehem et al., 2022; Frangou et al., 2022; Rutherford, Fraza, et al., 2022; Tamnes et al., 2017; Thambisetty et al., 2010). CT has been shown to more accurately reflect underlying pathophysiologic mechanisms than gray matter volume analysis (Clarkson et al., 2011; Hutton et al., 2009; Pereira et al., 2012; Zhao et al., 2022). However, these large data resources have expanded in scale via large, long-running longitudinal cohort studies. While the benefits of these large and unique cohorts are obvious, such studies impose particularly difficulties. For example, data must often be aggregated across multiple study centers, necessitating dealing with site effects and across developmental time scale, subjects are often scanned with different scanner hardware and/or software at successive timepoints. As a result, there is often little or no overlap in terms of age of participants and site effects in successive acquisition waves. Such non-trivial differences across sites, scanners, and timepoints have been difficult to account for statistically in analyses. Therefore, in addition to longitudinal data, novel methodological tools that map inter-individual differences are needed to generate new insights.

Normative modeling approaches have recently emerged as a tool for better understanding longitudinal developments with neuroimaging data (Marquand et al., 2019; Marquand, Rezek, et al., 2016). These approaches produce statistical inference at the individual level, without relying on strong assumptions about clustering of individuals or population structure (Antoniades et al., 2021; Cole, 2012; Marquand et al., 2019). Instead, symptoms in individual patients can be related to extreme deviation from the normative range (Fraza et al., 2021; Marquand, Wolfers, et al., 2016; Zabihi et al., 2019). This has shown the potential to detect morphological differences in patient populations which were not evident using standard techniques (Remiszewski et al., 2022). Additionally, a *Hierarchical Bayesian Regression* (HBR) approach to normative modeling has been shown to efficiently accommodate inter-site variation and to provide good computational scaling, which is useful when using large studies, longitudinal studies, or combining smaller studies together, that are acquired across multiple sites (Bayer et al., 2021; Kia et al., 2022; Rutherford, Fraza, et al., 2022). It also supports federated (*i*.*e*. decentralized) multi-site normative modeling to transfer previously trained models onto unseen sites, while benefiting from the training on the large reference datasets (Kia et al., 2022; Rutherford, Kia, et al., 2022). This is especially interesting given that in longitudinal studies running over several years, changes of scanner hardware, software and/or scan protocols are the norm rather than the exception, which generates a need to correct for the resulting scanner effects.

In this work, we provide a case study in using the transfer of prior knowledge about CT distributions from normative models derived from a large reference (e.g. lifespan) cohort to better estimate parameters on a smaller target (e.g. clinical) cohort. For this, we use longitudinal CT data from the Generation R study (Jaddoe et al., 2006; Kooijman et al., 2016; White et al., 2018), which contains data from children aged 6-17 years scanned in two different scanners, unseen by the reference models. The narrow age-range makes this study a good candidate for transfer learning in that it is necessary to transfer information learned from a large lifespan cohort to obtain precise estimates of the slope or trajectory of developmental effects across this narrow age range. This method provides important benefits: one, it allows meaningful comparison of individuals scanned on different scanners, while taking advantage of previous knowledge, built from large publicly available datasets to set informed hyperpriors: expected mean and variance of the distribution of samples for each ROIs. This, in turn, provide three benefits to the study, one providing more accurate predictions from the models thanks to the use of the mentioned informed priors; second this enables to reduce the ratio of training samples necessary to learn developmental trajectories for to the unseen sites, thereby enabling more participants to be allocated to the test set, and thus improving statistical power (Pan & Yang, 2010). Third, we will show that it provides a means to draw meaningful inferences within individuals across timepoints, even when follow-up scans are derived from a different scanner. This work also aims to offer some guidance on the methodology, *e*.*g*. providing empirical estimates of the number of samples required for the transfer of knowledge from previous learnings and choices in transfer configurations, *e*.*g*. factors included as batch effects. Finally, we provide a demonstration of the clinical utility of this approach by using it to understand inter-individual differences in brain morphology resulting from pre-term birth.

## 2. Methods

### 2.1 Normative modeling

We estimated normative models using *Hierarchical Bayesian Regression* (HBR) to predict cortical thickness from age, sex and scanner site, for each *region of interest* (ROI) using the PCNtoolkit python package, version 0.22 (Rutherford, Kia, et al., 2022).

#### 2.1.1 Reference models

We assembled a large reference cohort (Kia et al., 2022) containing n=40,435 (95%) healthy individuals to train the normative models before validating this model on n=2,548 controls and patients (5%, stratified by sites) from a collection of mostly publicly available MRI datasets across 77 sites and 42,983 participants: [ABIDE-1](http://fcon_1000.projects.nitrc.org/indi/abide/), [ADHD-200](https://fcon_1000.projects.nitrc.org/indi/adhd200/), [CAMCAM](https://www.cam-can.org/index.php?content=dataset), [PNC](https://www.med.upenn.edu/bbl/philadelphianeurodevelopmentalcohort.html), [CNP](https://www.openfmri.org/dataset/ds000030/), [HCP-Aging](https://www.humanconnectome.org/study/hcp-lifespan-aging), [HCP-Dev](https://www.humanconnectome.org/study/hcp-lifespan-development), [HCP-EP](https://www.humanconnectome.org/study/human-connectome-project-for-early-psychosis), [OASIS](https://www.oasis-brains.org/), [OPN](https://openneuro.org/), [IXI](https://brain-development.org/ixi-dataset/), [NKI-RS](http://fcon_1000.projects.nitrc.org/indi/enhanced/), [UKBB](https://www.ukbiobank.ac.uk/), [ABCD](https://abcdstudy.org/) and [CMI-HBN](http://fcon_1000.projects.nitrc.org/indi/cmi_healthy_brain_network/). The reference model is available on the PCNportal (https://pcnportal.dccn.nl/). Cortical thickness measures were obtained from FreeSurfer processing (versions 5.3 or 6.0), as referred in the publications associated with the datasets (Dale et al., 1999; Fischl et al., 1999, 2002; Fischl & Dale, 2000). Parcellation of the brain was made with the Destrieux atlas (Destrieux et al., 2010). One normative model was estimated per ROI. Linear HBR models were estimated using fixed effects of age and batch (*i*.*e*. random) effects for site and sex. In practice, this allows each site and sex to have different slopes, intercepts and variances. We included only data from the first visit when multiple visits were available (*i*.*e*. UKBB and ABCD). Any single missing individual ROI data (less than 0.1% of the samples per ROI) was imputed as the site and sex specific ROI mean.

Estimated reference models performed well according to accuracy metrics (explained variance: mean=0.44, SD=0.13, *standardized mean squared error* (SMSE): mean=0.55, SD=0.13, and *mean standardized log loss* (MSLL): mean=-0.37, SD=0.14). Outputs include hyperparameters defining the mean and variance of the site-specific mean effects and variance, estimated during the training over the collection of datasets. This can be used as informed priors when adapting the normative models to unseen target sites. These hyperparameters are adapted to the unseen site using a holdout subset of the target dataset, *i*.*e*. the adaptation set. This allows to reduce the number of samples used for adaptation while retaining a low variance of the estimations.

#### 2.1.2 Target cohort

As target cohort, 8,523 T_1_-weighted MRI scans from the population-based longitudinal Generation R study (Jaddoe et al., 2006; Kooijman et al., 2016) were used. In short, the Generation R study is a prospective cohort study from fetal life until adulthood that is designed to find early markers for typical and atypical development, growth, and health. Almost 10,000 pregnant women living in Rotterdam, the Netherlands, were enrolled in the study between 2002 and 2006. Data from the children and caregivers was collected at several time points and written informed consent and/or assent was obtained from all participants. The imaging protocol and quality assessment is extensively described by White, Muetzel and colleagues (White et al., 2018). MRI scans were acquired in 3 waves using 2 different scanners, making the cohort an ideal validation set to investigate the transfer of hyperparameters from a reference dataset to an unseen target set. In longitudinal studies running over several years, changes of scanner hardware, software and/or scan protocols are often inevitable, which generates a need to correct for the resulting scanner effects. In the first wave, 1,033 participants (484 female, age range: [6-10]) were imaged with a 3T MR750 Discovery MRI scanner, while in the second (n=3,920, 1,977 female, age range: [9-12]) and third wave (n=3,570, 1,866 female, age range: [13-17]) a 3T MR750w Discovery scanner (General Electric, Milwaukee, WI, USA) was used. After exclusion of scans with incidental findings (n=58), braces (n=1067), and low-quality visual inspection ratings of FreeSurfer reconstructions (n=2067), a total of 6,285 scans were included in the target dataset. Figure 1 shows a histogram of age and scanner distributions in the target dataset.

**Figure 1.**
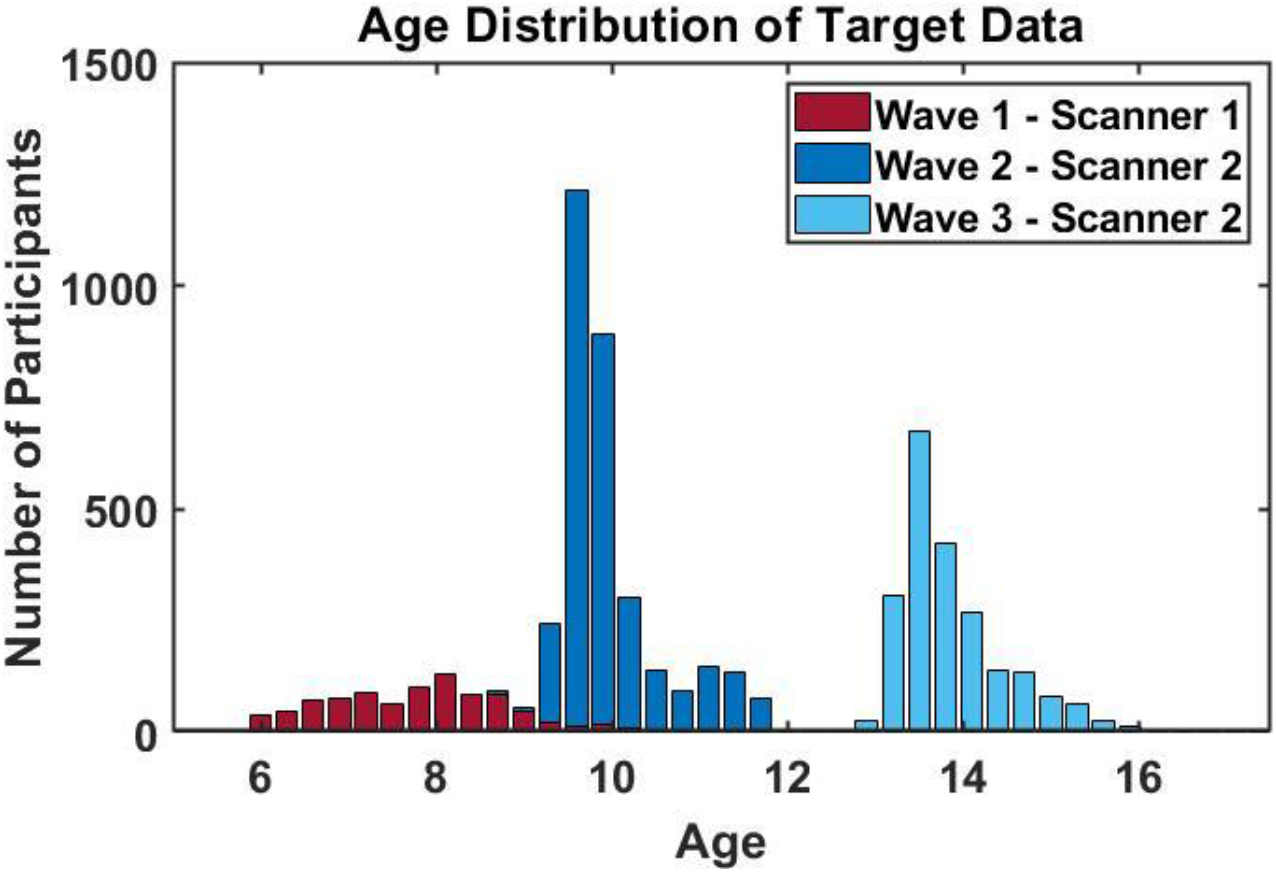
Histogram of the scanning waves and age distributions in the Generation R target dataset.

### 2.2 Transfer of hyperparameters from reference models to target cohort

By making use of the Generation R study cohort, we set out to show the advantage of transferring the hyperparameters to an unseen site by 1) determining the optimal number of samples needed for adaptation to the target cohort, 2) validating the recalibration of data to the target cohort and successful removal of site-effects by comparing raw and scanner corrected values, and 3) illustrating the utility of site-corrected deviations scores to uncover changes in morphology between groups and individuals. In the following, these three aims are described in more detail.

#### 2.2.1 Optimal sample size for parameter adaptation

In order to determine the optimal number of samples in the adaptation set we leveraged the large amount of data available in the Generation R dataset. As described above, to prevent bias, held-out data should be used for adapting the parameters of the normative model to the target cohort (see Kia et al., 2020, 2022 for details). The number of scans in the adaptation set was varied ranging from 5 to 300 scans and model metrics (explained variance, SMSE, MSLL) of the subsequent models was calculated for each sample size. The resulting information is particularly useful for small imaging cohorts, since cohorts with smaller sample sizes can employ the current approach to boost power by making use of the hyperpriors inferred from large data. Yet, this is only viable if the samples needed to recalibrate the models can be kept to an optimal minimum.

#### 2.2.2 Validation of adaptation

Additionally, two aspects of the Generation R study design make the cohort an ideal target set to validate the successful recalibration of the normative models to an unseen site. First, scans of participants that have repeated measurements over all 3 scanning time points are present. Uncorrected CT values of a participant with scans across all 3 measurement waves (and therefore across both scanners) show heterogeneity over time points that is partly due to changes over time and partly due to confounding site-effects. After successful recalibration of the normative model, we expect resulting z-scores, which are in principle free of site-effects, of the same participant to be in a similar range while raw values will differ. Second, there is an overlapping age range (8,6 - 10,7 years of age), in which scans from both scanners were obtained (Fig. 1). Z-scores of participants from wave 1 (scanner 1), that fall in the overlapping age range of wave 1 and wave 2 should be distributed similarly after recalibration as z-scores in the same age range of wave 2 (scanner 2) while raw, uncorrected values differ due to scanner effects. Therefore, scans of participants with measurements at all 3 scanning time points (n=1,317) and scans from the first imaging wave that fall in the overlapping age range (n=211) were withheld from the adaptation set that was used to recalibrate the reference normative model to the new unseen site. As outlined above, these scans hold valuable information that will be used to determine the successful calibration of the models by comparing raw CT before adaptation and corrected estimates after adaptation.

#### 2.2.3 Clinical application of normative estimates

Lastly, we used the resulting site-effect free estimates to illustrate their potential to uncover morphological deviations in clinical cohorts by contrasting estimates in CT per ROI between participants in the Generation R cohort born pre-term (gestational age < 37 weeks, n=339) and children born at term (n=5,646). Pre-term birth interrupts a vulnerable period for brain development, as processes such as synaptogenesis, axonal growth, and neuronal migration, take place during the third semester (Volpe, 2009). Therefore, deviation scores from the normative models can for instance be used to explore the variability in CT within children born pre-term, but also to find ROIs that differ between children born pre-term and at term. Notably, these deviation scores are free of site-effects and therefore especially suited for longitudinal MRI designs, as it is the case with the Generation R study.

## 3. Results

### 3.1 Transfer results

#### 3.1.1 Optimal number of samples for parameter adaptation

We first determined the optimal number of subjects needed in the adaptation set. Figure 2 shows evaluation metrics for each ROI as the sample size of the adaptation set increases. Performance of the model reaches a plateau around 100 subjects. We thus adapted the initial reference models to the unseen sites of the Generation R study on n=300 (4.8%) (n=100 for scanner 1 in wave 1; n=200 for scanner 2 in wave 2 and 3) and tested the models on the remaining participants (n=5,985; n=813 for scanner 1 in wave 1, n=5,172 for scanner 2 in wave 2 and 3). Subjects from wave 1 and 3 were sampled randomly, whereas subjects from wave 2 were sampled pseudo-randomly to ensure a uniform cover of the full range of the narrow and highly peaked age distribution in this wave (Fig. 1). While model performance reached a performance ceiling at approximately 100 scans per scanner/wave in the adaptation sample, only slight concessions in model performance are present as adaptation sample size decreases to only 25 scans.

**Figure 2.:**
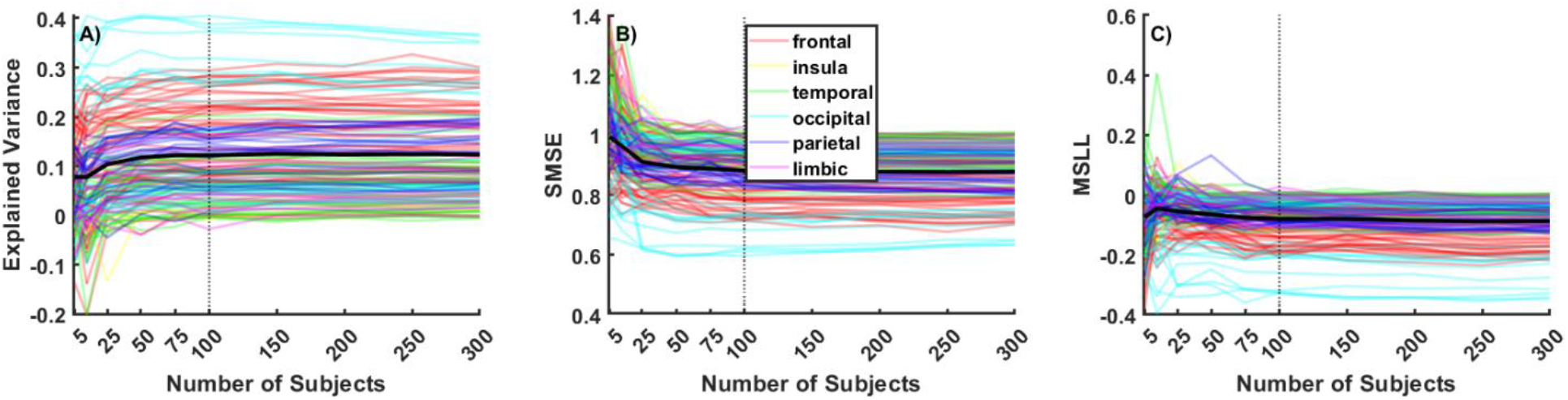
Comparison of model performance as the number of subjects in adaptation set increases. Colored lines show evaluation metrics per ROI, color coded according to cerebral area. The black line illustrates the mean across all ROIs. Model performance reaches a plateau at approximately 100 scans per wave in the adaptation sample (vertical dotted line).

#### 3.1.2 Adaptation settings

We furthermore tested different adaptation settings. Despite the fact that scans in wave 2 and 3 of the target cohort were acquired on the same scanner, we compared adaptation settings treating waves 2 and 3 as the same but also as different sites. When treated as same sites, we found a slight bias for higher deviation scores (z-scores) when running the adaptation to a wave 2+3 test set with wave 3 subjects compared to wave 2 subjects only, in particular for frontal ROIs. The effect of the different adaptation settings on all ROIs is shown in Supplementary Figure 1 and is explicitly illustrated in an example ROI in Figure 3. Panel A shows that the model is more successful in reparametrizing the raw data to centiles when each time point of measurement is handled as a separate site-effect. Possible sources for such effects might stem from changes in scanner software, changes in image quality with age (i.e. motion artifacts), or sample variability. In our target cohort, scanner software was upgraded after the first 370 scans of wave 2 but was otherwise identical in wave 2 and wave 3. However, age-related improvements in images quality are frequently reported in the literature and quality assurance, measured as topological defects in the surface reconstruction for FreeSurfer processed MRI data (https://github.com/Deep-MI/qatools-python), does show improvements in image quality with age across the three waves (Mean_wave 1_=229.06, SD_wave 1_=98.56; Mean_wave 2_=213.89, SD_wave 2_=67.15; Mean_wave 3_=166.97, SD_wave 3_=67.15).

**Figure 3.**
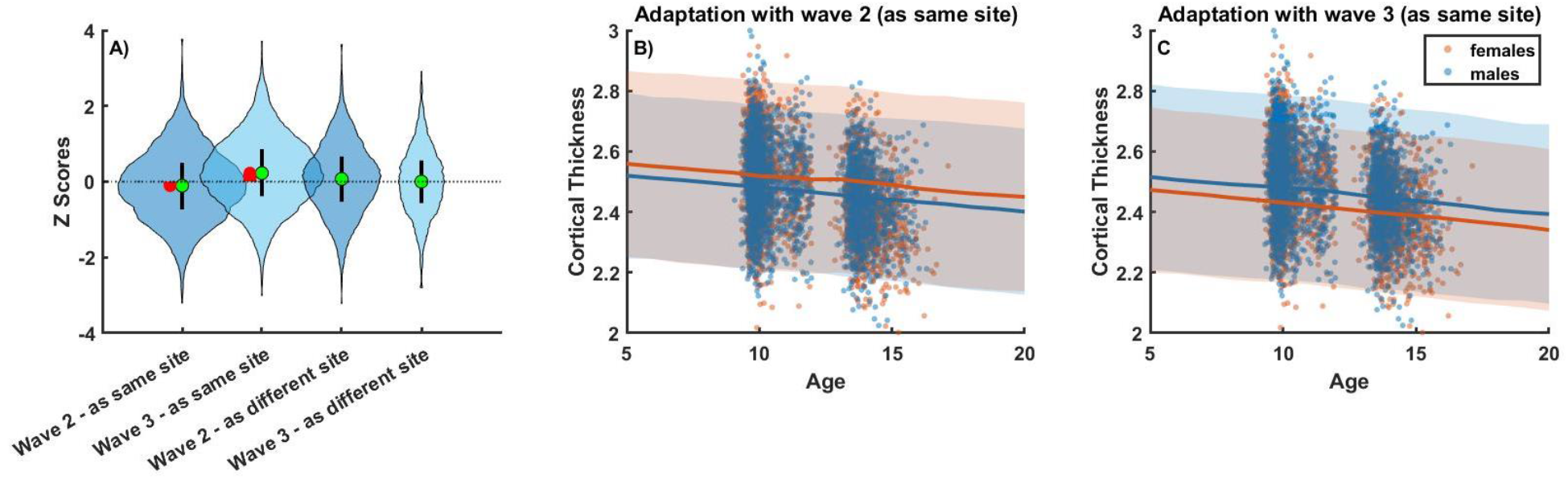
Effects of different recalibration configurations on the target cohort illustrated in an example ROI (inferior frontal sulcus). Panel A) shows z-score distributions when measurement waves 2 and 3 of the Generation R target cohort are treated as the same (same scanner) or different sites in a frontal example ROI, the inferior frontal sulcus. Median and interquartile range are represented by green dots and black bars, respectively. For each measurement wave, we would expect the median z-score to be around 0. However, this is not the case if measurement wave 2 and 3 are treated as the same site. The difference from 0 is indicated by red bars. By examining the CT trajectories in panel B) and C), we see that this might be due to a misestimation of mean and variance in females when both waves are treated as the same site.

#### 3.1.3. Adaptation Validation

After choosing for a adaptation setting treating the three measurement waves as different batch effects, we validated the success of the adaptation of the reference model to the target cohort by examining the differences between raw CT values and corrected deviations (z-scores) after transfer of the subjects which were withheld from the adaptation sets (Fig. 4). Scans of participants with repeated measurements at all imaging waves (a random sample of ten participants is depicted by green lines) show a decline over time in raw CT. As expected, thinning of the cortex can be observed with age, however, the raw CT values are confounded by noise stemming from site-effects of the different measurement waves. In the resulting z-scores of the withheld subjects, these site-effects are removed as demonstrated by stable deviations from the normative model within a participant (Fig. 4B). The same holds true for the withheld subjects from measurement wave 1 that fall in the overlapping age range (8,6 - 10,7 years of age) of wave 1 (scanner 1) and 2 (scanner 2). While raw CT values in the overlapping age range vary vastly between the two measurement waves (*t*(2874)=13.4, *p*<0.001), with a tendency of higher values in measurement wave 1 compared to wave 2 (Fig. 4C), this difference is slightly reduced when correcting for sex (Fig. 4D) (*t*(2874)=11.4, *p*<0.001) and practically absent in the sex- and additionally site-effect corrected z-scores (Fig. 4E) (*t*(2874)=1.0, *p*=0.324). Therefore, we can meaningfully compare individuals on the basis of z-scores, bearing in mind that the z-scores are defined with respect to a lifespan based normative model.

**Figure 4:**
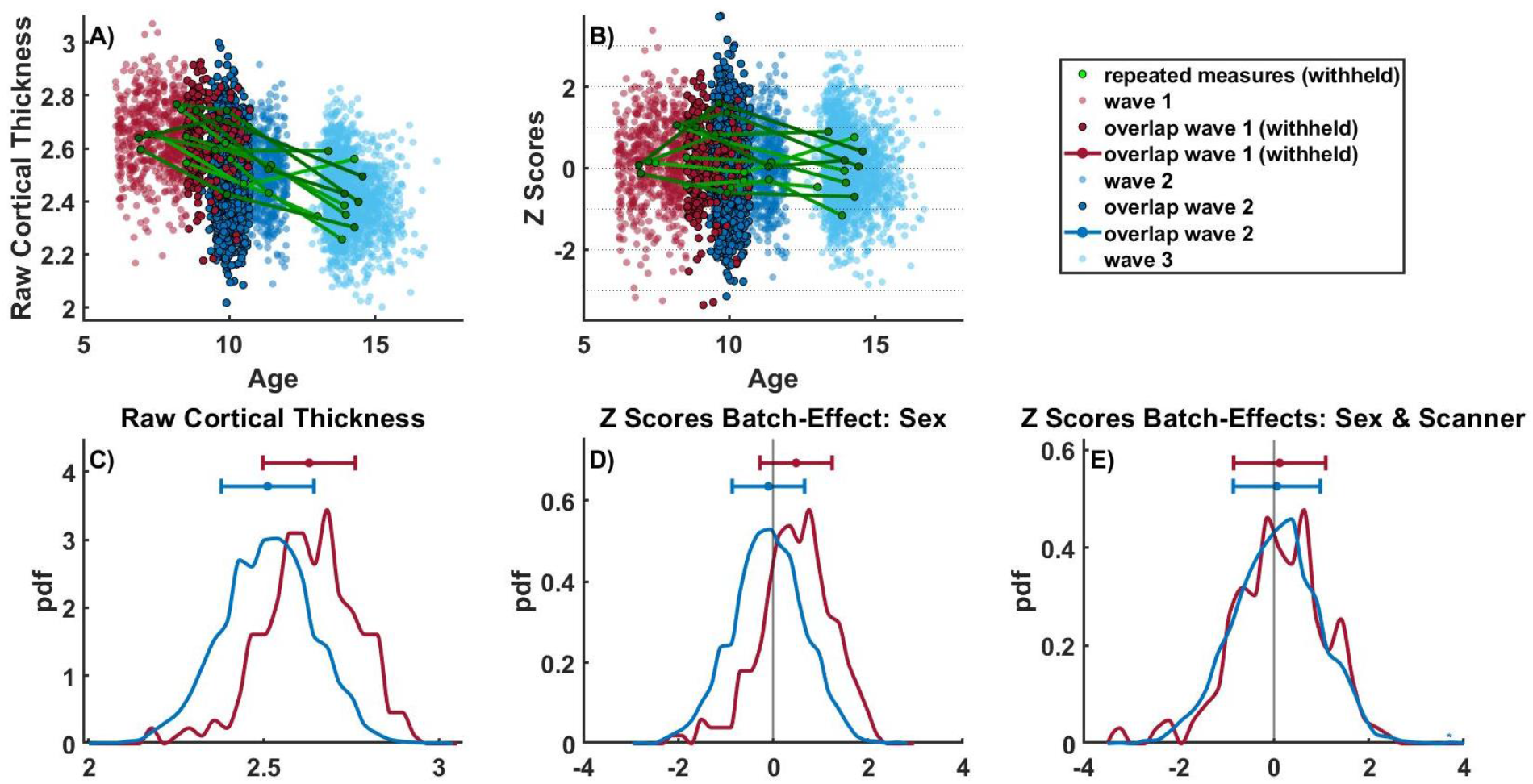
Validation of transferring the reference normative model to the target cohort using two groups of subjects that were withheld from the adaptation set: 1) subjects with repeated measurements at all 3 imaging waves (random sample of ten participants depicted by green lines, panels A and B; 2) subjects from imaging waves 1 and 2 that fall in the overlapping age range of both scanners [8,6 - 10,7] (depicted by darker shaded red and blue dots and lines, panels A-E). Panel A and C show raw cortical thickness values. Panels B, D, and E show sex-effect (panel D) or sex- and site-effect corrected z-scores of the same participants (panels B&E). For consistency, the same ROI (inferior frontal sulcus) as in the previous figures is illustrated.

### 3.2 Relating site-effect corrected z-scores to gestational age

To illustrate the usefulness of the resulting models, we compared extreme deviations, acquired at the level of individuals, between children born pre-term and children born at term in the target cohort. Percentages of individuals with an extreme z-score (larger/smaller than 2) per ROI are shown in Figure 5. In the children born at term, we find approximately 2.5% of children with extreme negative and extreme positive z-scores respectively across ROIs. Exceptions are primarily smaller ROIs (sulcus intermedius primus (left & right), posterior ramus of the lateral sulcus (left), anterior transverse collateral sulcus (right), orbital sulcus (right)) where areas with thicker cortices than expected can be observed. Importantly, extreme deviations are much more prevalent with children born pre-term with the most pronounced extreme positive deviations (thicker cortex than expected) found in the left pericallosal sulcus and lateral aspect of the superior temporal gyrus, as well as in the anterior part of cingulate gyrus and sulcus (ACC) of both hemispheres. The most striking extreme negative deviations (thinner cortex than expected) can be seen on the left hemisphere in the superior and inferior temporal sulcus, lingual sulcus, superior part of the precentral sulcus, supramarginal gyrus, and on the right hemisphere in the superior and inferior part of the precentral sulcus, superior frontal sulcus, angular gyrus, precentral gyrus and the precuneus.

**Figure 5.:**
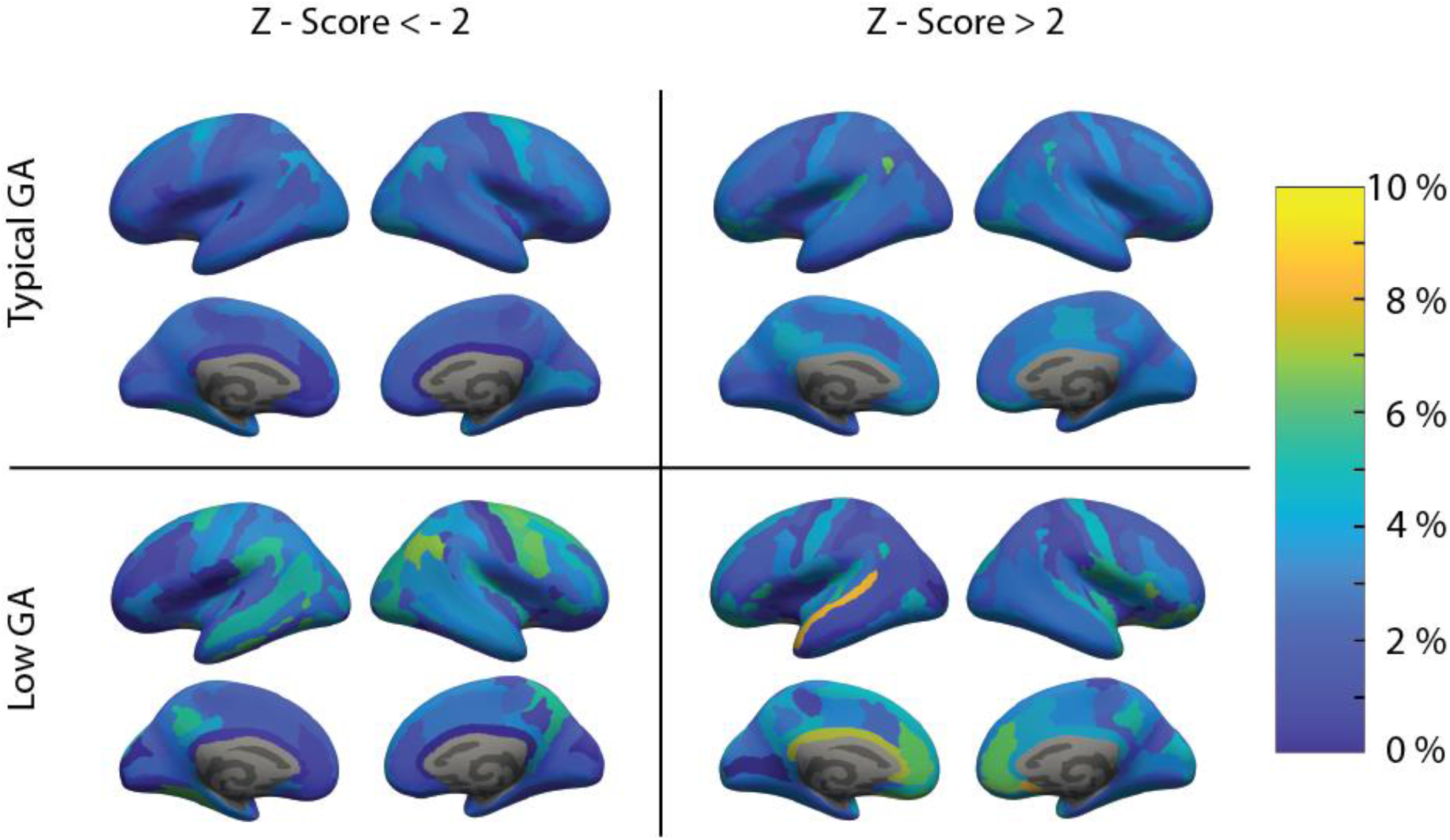
Differences in site-effect corrected z-scores between children born pre-term (low gestational age (GA)) and children born term (typical gestational age (GA)). On the left side, extreme negative deviations (cortex thinner than expected) are illustrated. On the right, extreme positive deviations (cortex thicker than expected) are shown.

These regions are consistent with previous findings on CT differences in adolescents born pre-term. Pronounced cortical thinning has been found persistently in areas surrounding the central sulcus and temporal lobes (Martinussen et al., 2005; Nagy et al., 2011; Zubiaurre-Elorza et al., 2012) as well as thicker cortices in frontal regions surrounding the anterior cingulate cortex (Bjuland et al., 2013). The current approach has been shown to capture structural deviations better than case-control studies as they are more sensitive to individual heterogeneity (Remiszewski et al., 2022). It also offers improved insights in longitudinal cohorts, as these deviation scores are not cofounded by site-effects.

## 4. Discussion

In this study, we used information from normative models that were initially trained on a large number of samples, scanned over 77 sites, as prior knowledge for the parameters of the CT distributions when adapting these models to the two scanners of the longitudinal Generation R study.

We report three main findings: first, transfer learning is successful and allows for meaningful comparisons between individuals from different scanners, and sexes, as previously reported (Kia et al., 2022). Second, we quantified the number of samples in the transfer set needed to obtain good performance metrics on the test set and show that relatively few samples are sufficient for good performance (approximately n=25). This provides the added benefit of improving the statistical power of statistical analyses on the resulting larger test set. While we used 100 samples per measurement wave in the adaptation site, slightly smaller adaptation samples decreased the evaluation metrics only marginally. Third, we show that the deviations from these normative models are meaningful in that they are altered in a highly individualized manner in individuals born pre-term.

Our results support the finding that normative models capture the general trend of decreasing cortical thickness with age, as reported in previous studies (Bethlehem et al., 2022; Frangou et al., 2022; Rutherford, Fraza, et al., 2022; Tamnes et al., 2017; Thambisetty et al., 2010). Interestingly, we found that the model performed better when each measurement wave of the transfer cohort was treated as a separate site-effect, even though two of three waves were acquired on the same scanner. This could be due to sample variability or a misestimation of parameters in the female cohort, possibly linked to the fact that scan quality tends to improve with age. For future studies, it may be useful to treat distinct measurement intervals as separate batch-effects, resulting in a factorial design of sex x scanners x waves, even if the scanner setup has not changed, to produce more precise models. Our recommendations might differ for longer timescales, non-linear or non-Gaussian lifespan trajectories, which usually requires more data (de Boer et al., 2022). However, the methods we introduce can be used to determine the optimal number of subjects for such cases.

The successful validation of the use of transfer learning with normative models opens the door for further investigations exploring the relationship between deviation scores and various phenotypes. Individual-level deviations, as obtained through normative models, have been shown to provide stronger effects than typical case-control studies using uncorrected raw measurements (Rutherford, Fraza, et al., 2022) and are therefore particularly suitable for exploring and investigating individual differences within and across datasets. The used federated learning framework makes it possible to use the models presented in this work as informed priors (models are available online via PCNportal [https://pcnportal.dccn.nl/] to investigate CT in smaller and/or clinical cohorts.

In this study, we provide an example how normative models can be used to investigate clinical phenotypes, by investigating the relation between extreme deviations scores and the gestational age at birth, that is between children born at-term and pre-term. While children born at-term show an expected distribution of approximately 2.5% with z-scores higher than 2 or lower than -2 respectively, children born pre-term are more likely to have extreme deviations in specific ROIs, which are consistent with previous literature showing pronounced differences in particular in frontal and temporal cortices. While we show a comparison between groups, the current approach does not require clustering of individuals into groups but instead can be used to make inferences about heterogeneity within clinical groups as well as about deviations on an individual level.

### 4.1. Limitations and future directions

Although we demonstrate that evaluation metrics level off after 100 scans in the adaptation set, as few as 25 scans can still lead to effective transfer of knowledge. However, this limitation prevents small cohorts from utilizing the current approach due to the fact that the median sample size of neuroimaging studies typically includes 25 participants (Marek et al., 2022).

Furthermore, our work estimates normative models on a single ROI, thereby neglecting any spatial interdependencies between brain regions. Moreover, other image-derived phenotypes, such as the cerebellum, could also be considered.

## 5. Conclusion

Using longitudinal cortical thickness data from the Generation R study on children aged 6 to 17 years old, we present an application of transfer learning of large-scale normative models which produce good performance metrics with even a limited size of adaptation set. The resulting deviation scores per age and ROIs, allow for meaningful comparison inter sites and inter sex.

Using these obtained deviation scores, we were able to show specifically localized differences in cortical thickness between children born pre-term and children born at-term.

## Supporting information

Supplementary Figure 1.: Effects of different recalibration configurations

