## Supplementary Figure 1.: Effects of different recalibration configurations for "Estimating cortical thickness trajectories in children across different scanners using transfer learning from normative models"

Supplementary Figure 1.: Effects of different recalibration configurations on the target cohort illustrated in all ROIs of the Destrieux parcellation. ROIs are grouped in frontal, parietal, temporal, insular and limbic, and occipital lobes.

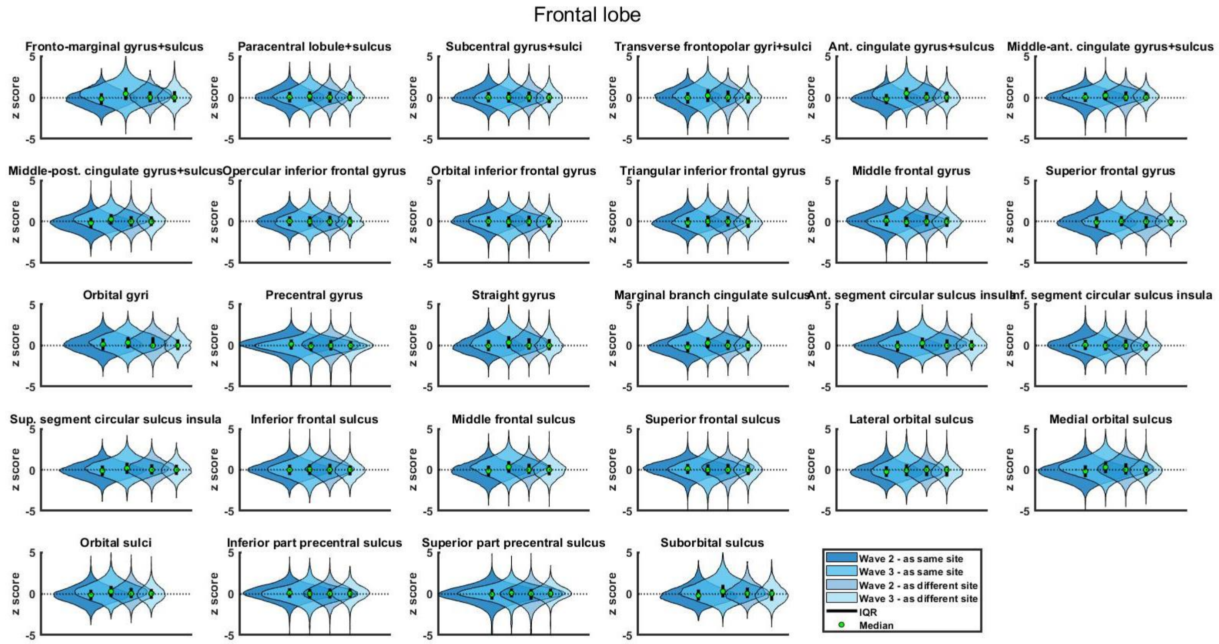


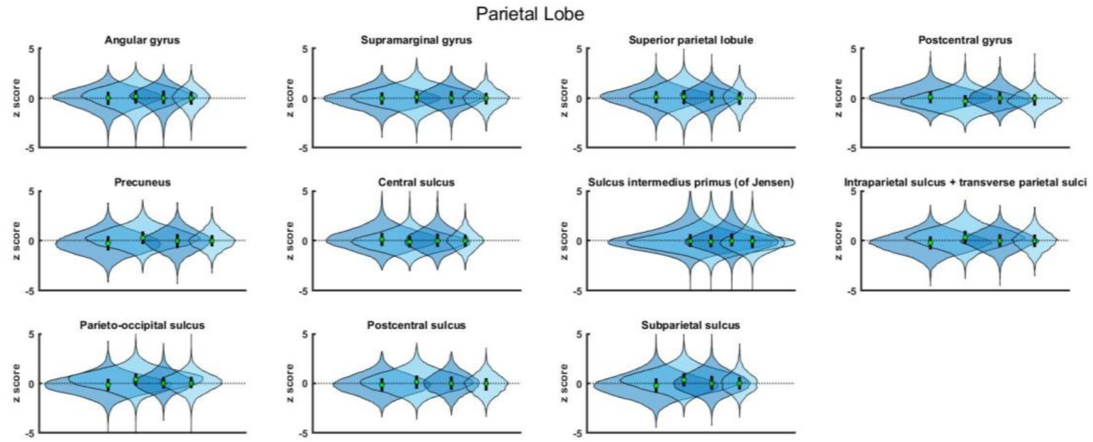

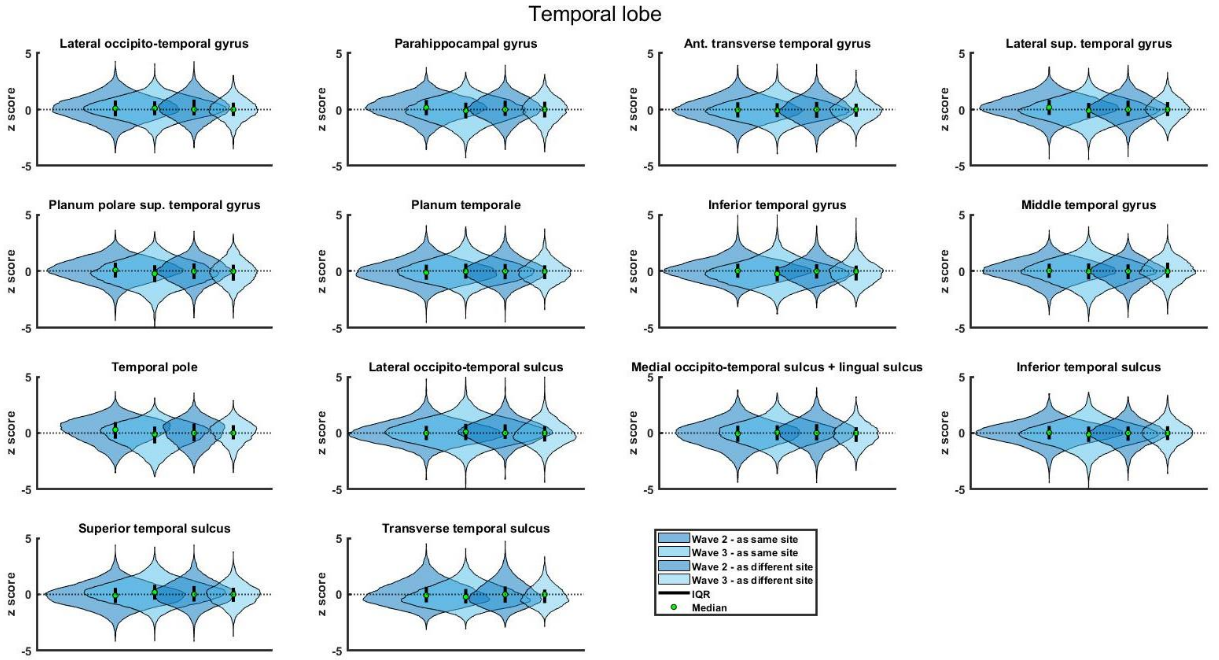
Temporal lobe
Lateral occipito-temporal gyrus
Parahippocampal gyrus
Ant. transverse temporal gyrus
Lateral sup. temporal gyrus
Planum polare sup. temporal gyrus
Planum temporale
Inferior temporal gyrus
Middle temporal gyrus
Temporal pole
Lateral occipito-temporal sulcus
Medial occipito-temporal sulcus + lingual sulcus
Inferior temporal sulcus
Superior temporal sulcus
Transverse temporal sulcus
z score
5
0
-5
Wave 2 - as same site
Wave 3 - as same site
Wave 2 - as different site
Wave 3 - as different site
IQR
Median

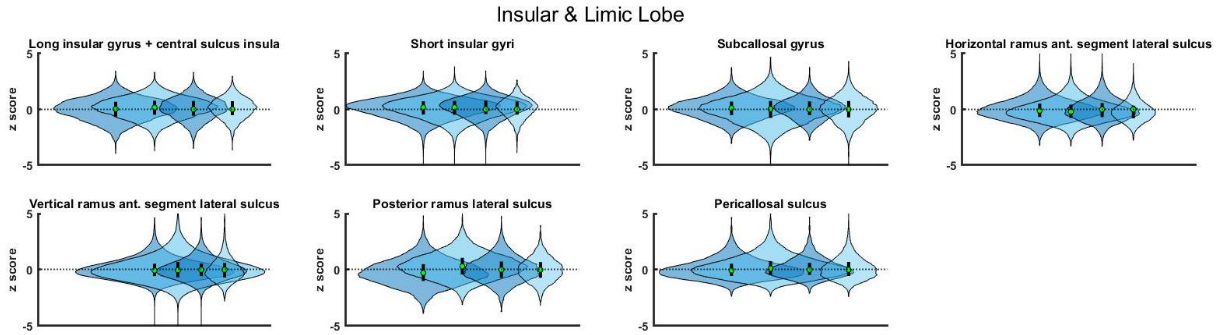
Insular & Limic Lobe
Long insular gyrus + central sulcus insula
Short insular gyri
Subcallosal gyrus
Horizontal ramus ant. segment lateral sulcus
Vertical ramus ant. segment lateral sulcus
Posterior ramus lateral sulcus
Pericallosal sulcus
z score
5
0
-5
